# Age-related dedifferentiation and hyperdifferentiation of perceptual and mnemonic representations

**DOI:** 10.1101/2020.06.15.151217

**Authors:** Lifu Deng, Simon W. Davis, Zachary A. Monge, Erik A. Wing, Benjamin R. Geib, Alex Raghunandan, Roberto Cabeza

## Abstract

Preliminary evidence indicates that occipito-temporal activation patterns for different visual stimuli are less distinct in older (OAs) than younger (YAs) adults, suggesting a dedifferentiation of visual representations with aging. Yet, it is unclear if this deficit (1) affects only sensory or also categorical aspects of representations during visual perception (perceptual representations), and (2) affects only perceptual or also mnemonic representations. To investigate these issues, we fMRI-scanned YAs and OAs viewing and then remembering visual scenes. First, using representational similarity analyses, we distinguished sensory vs. categorical features of perceptual representations. We found that, compared to YAs, sensory features in early visual cortex were less differentiated in OAs (i.e., age-related dedifferentiation), replicating previous research, whereas categorical features in anterior temporal lobe (ATL) were more differentiated in OAs. This is, to our knowledge, the first report of an *age-related hyperdifferentiation*. Second, we assessed the quality of mnemonic representations by measuring encoding-retrieval similarity (ERS) in activation patterns. We found that aging impaired mnemonic representations in early visual cortex and hippocampus but enhanced mnemonic representations in ATL. Thus, both perceptual and mnemonic representations in ATL were enhanced by aging. In sum, our findings suggest that aging impairs visual and mnemonic representations in posterior brain regions but enhances them in anterior regions.

## 1. Introduction

As we age, the anatomy and physiology of our brain declines, impairing cognitive abilities such as perception and memory (Grady, 2012; Grady, 2008). Most prior functional MRI (fMRI) studies investigating the neural bases of these impairments have focused primarily on *processes* (operations performed on information) and only rarely examined age effects on *representations* (the nature of the information processed) (Cowell et al., 2019). In fMRI studies, differences in the mean activity level (univariate measure) of a group of voxels is assumed to reflect differences in processes, whereas differences in the spatial distribution (pattern) of activity (multivariate measure) within a group of voxels is assumed to reflect differences in representations (Haxby et al., 2001; Norman et al., 2006; Kreigeskorte et al., 2008). There has been accumulating evidence that activation patterns elicited by different types of visual stimuli (faces, places, etc.) are less distinct in older adults (OAs) than in younger adults (YAs; Park et al., 2004; Chee et al., 2006; Payer et al., 2006; Voss et al., 2008; Goh et al., 2010; Bowman et al., 2019; Koen et al., 2019; Koen and Rugg, 2019; Koen et al., 2020). This phenomenon, known as *age-related neural dedifferentiation*, suggests that aging impairs the quality of representations during visual perception (*perceptual representations*). Yet, two fundamental questions remain unanswered: (1) what aspects of perceptual representations are impaired by aging? and (2) are age-related deficits in perceptual representations associated with deficits in visual memory traces (*mnemonic representations*)? The current study investigates these two critical questions.

### 1. What aspects of the perceptual representations are impaired by aging?

This is a critical issue because perceptual representations consist of multiple features, which are processed in different brain regions and are affected differentially by aging. In particular, it is well established that sensory features of visual representations are processed primarily in early visual cortex and that these features form conjunctions of categorical features in more anterior ventral pathway regions, such as the anterior temporal lobe (ATL; Bussey et al., 2005; Clarke and Tyler, 2014). The sensory-to-categorical feature distinction is relevant to aging because OAs tend to be impaired in perceptual processes (Park et al., 2004; Chee et al., 2006; Goh et al., 2010; Bowman et al., 2019; Koen et al., 2019; Koen and Rugg, 2019; Koen et al., 2020). However, their abilities to process categorical and conceptual information are relatively preserved (Cherry et al., 2012; Mohanty et al., 2016; Monge and Madden, 2016; Owsley, 2011), suggesting that age-related perceptual impariment could be driven more by the deficit in processing lower-level sensory information. Thus, we hypothesized that *age-related visual dedifferentiation impairs sensory features of perceptual representations in early visual cortex but not categorical features in the ATL* (**Hypothesis 1**). Given than some aspects of categorical-related processing are actually better in OAs than YAs (Long and Shaw, 2000; Park et al., 2002), an intriguing possibility is that categorical features in the ATL could be enhanced by aging.

We investigated Hypothesis 1 using representational similarity analyses (Kriegeskorte and Kievit, 2013; Kriegeskorte et al., 2008), in which the *sensory* and *categorical* similarities between stimuli are coded by two separate *stimuli models*. In the *sensory model*, pair-wise stimuli similarity is based on sensory visual features, such as shape (e.g., gun ≈ hair dryer), whereas in the *categorical model*, it is based on categorical features (e.g., gun ≈ sword). The pair-wise stimuli similarity coded by the model is then correlated with pair-wise similarity in fMRI activation patterns (representations) for the same set of stimuli. The resulting *model-brain fit* (2^nd^ order correlation) identifies brain regions that process and/or store representations emphasizing sensory and/or categorical visual features (**Fig. 1a**). As stimuli models code the differences between images based upon a selective feature of interest (e.g., sensory, categorical), this model-brain fit may be operationalized as a measure of differentiation, where higher values in a brain region indicate greater neural differentiation.

**Figure 1:**
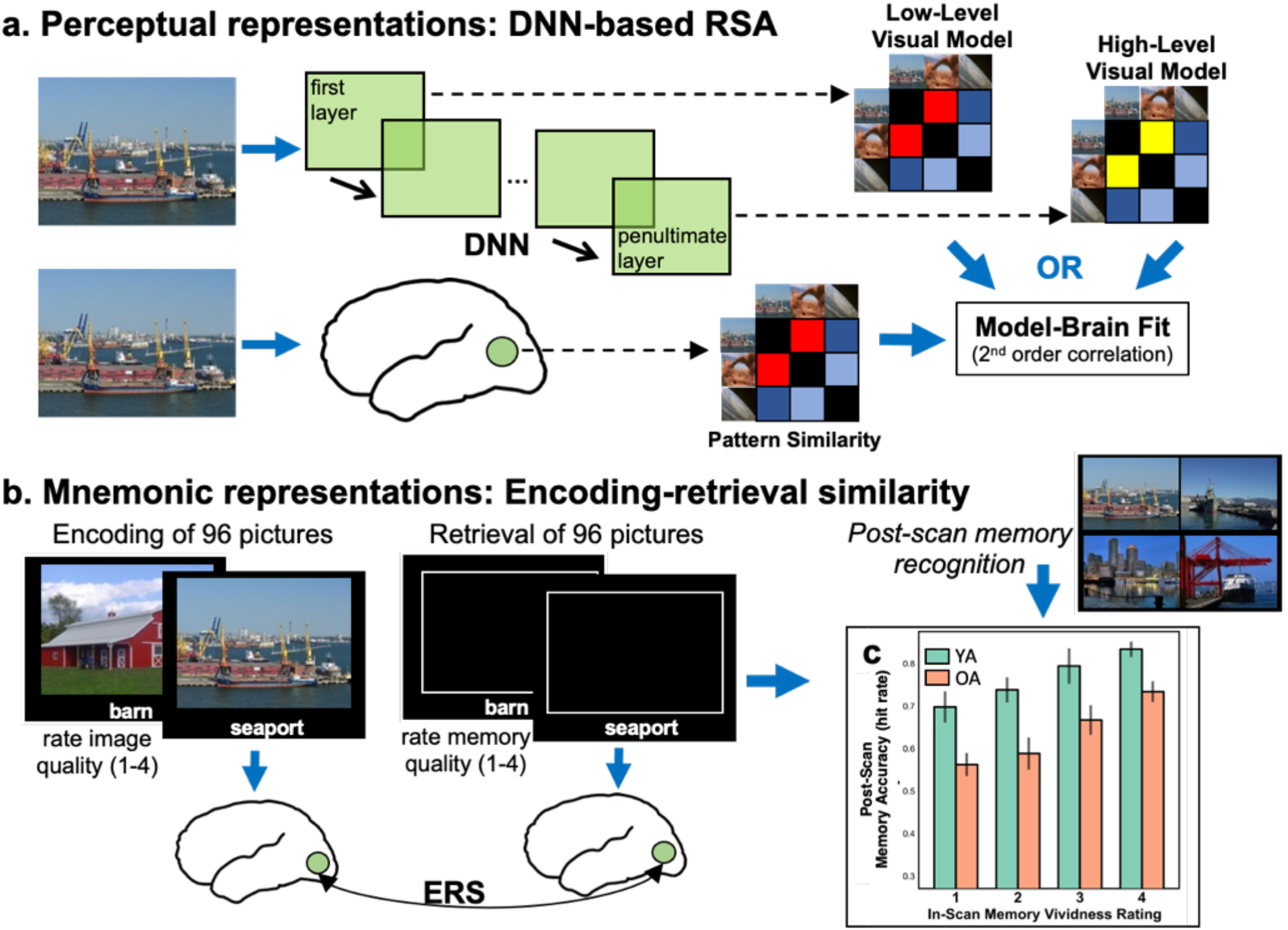
Perceptual and mnemonic representations analysis approach. **Panel a** shows a model of the analysis combining representational similarity analysis and DNN layers. To create models from DNN layers, the image stimuli are submitted to the pre-trained DNN and at each layer of interest, the “activation” values are extracted. Here, our layers of interest were the first and penultimate layers. For each image and layer of interest, the vectors of activation values are correlated with each other resulting in a stimuli model for each layer of interest. The stimuli models may then be used to predict brain activation patterns (model-brain fit) within ‘searchlight’ volumes (green circle). **Panel b** shows our paradigm in which participants, while undergoing fMRI scanning, studied images (while rating the quality of the images) and later retrieved their memories of the scenes (while rating the vividness of their memories). We quantified the similarity of encoding to retrieval representations by calculating ERS. **Panel c** shows for each in-scan vividness rating value, the corresponding post-scan memory accuracy (hit rate). Error bars represent the standard error of the mean. ERS = encoding-retrieval similarity; DNN = deep convolutional neural network; fMRI = functional MRI; OA = older adult; RSA = representational similarity analysis; YA = younger adult.

In the current study, our stimuli models were derived from *deep convolutional neural networks* (DNNs; Krizhevsky et al., 2012; LeCun et al., 2015). DNNs consist of layers of convolutional filters and can be trained to classify images into categories with a high level of accuracy. During training, DNNs “learn” convolutional filters in service of classification in the final layer, where filters from early layers predominately detect sensory features and while later layers organize items by their categorical features (Zeiler and Fergus, 2014). Previous work has demonstrated that progressive layers of typical DNNs (AlexNet, VGG16, etc.) may help to model visual representations along the ventral visual pathway, where early layers map brain activation patterns predominately in early visual cortex and late layers map brain activation patterns in more anterior ventral visual pathway regions (Güçlü and van Gerven, 2015; Khaligh-Razavi and Kriegeskorte, 2014; Kriegeskorte, 2015; Leeds et al., 2013; Wen et al., 2017; Davis et al., 2020). Thus, a principle advantage of DNNs is that they afford the ability to model a continuum of low-level sensory features to more semantically meaningful categorical features. These models are especially helpful in addressing the complex visual arrays typical of most scenes (as opposed to single objects, Josephs and Konkle, 2019), especially with scenes where their categories are less clear. DNNs allow for this hierarchy to be investigated within a single framework. Furthermore, DNNs outperform traditional theoretical models of the ventral visual pathway (e.g., HMAX, object-based models; Cadieu et al., 2014; Groen et al., 2018) in their capacity to identify specific objects with the appropriate category- or basic-level label. Therefore, a DNN is an ideal model to investigate this sensory-to-categorical feature distinction. Here, we used a pre-trained 16-layer DNN, the VGG16 (Simonyan and Zisserman, 2014), which was successfully trained to classify 1.8 million scenes into 365 categories (Zhou et al., 2017). The first hidden layer generated a *sensory model*, since this model is derived from a layer that detects sensory features, and the penultimate layer, a *categorical model*, since this model is derived from the layer before the images are explicitly categorized into the trained categories (Bankson et al., 2018; Devereux et al., 2018; Groen et al., 2018).

Given the posterior-anterior organization of the ventral visual pathway, we expected that the sensory model would correlate with activation patterns in early visual cortex, and the categorical model, with activation patterns in more anterior temporal regions such as the ATL. On the basis of Hypothesis 1, we predicted that, compared to YAs, (1) the correlation between brain activation patterns and the sensory model (i.e., model-brain fit) would be reduced in OAs, whereas (2) the correlation between brain activation patterns and the categorical model would be spared (or even enhanced) in OAs.

### 2. Are age-related deficits in perceptual representations associated with deficits in mnemonic representations?

Age-related sensory and cognitive deficits are strongly related to each other (Baltes and Lindenberger, 1997; Lindenberger and Baltes, 1994), possibly because sensory deficits cascade through the cognitive system impairing downstream cognitive processes (Monge and Madden, 2016). Consistent with this idea, degrading stimuli (i.e., mimicking sensory impairment) by adding noise yields cognitive deficits in YAs that resemble cognitive deficits in OAs (Gilmore et al., 2006; Monge and Madden, 2016; Murphy et al., 2000; Pichora-Fuller et al., 1995). In contrast, spared categorical processing in OAs may explain why age-related memory deficits are attenuated for semantically rich stimuli (Kausler, 1994; Naveh-Benjamin, 2000). Thus, the sensory to categorical dissociation we postulated for perceptual representations (Hypothesis 1) is likely to apply also to mnemonic representations.

We operationalize “perceptual representations” as the model-brain fit during the initial image presentation during the encoding phase of our study. Two types of perceptual representations are investigated, sensory representations and categorical representations. In contrast, we operationalize “mnemonic representations” in terms of the reactivation of the encoded activation during retrieval, as measured by the similarity between encoding and retrieval activation patterns or *encoding-retrieval similarity* (ERS, Fig. 1b). In the case of mnemonic representations, however, the age-related deficit is likely to affect not only visual cortex but also downstream memory-binding regions, such as the hippocampus. The hippocampus shows strong evidence of reactivation during retrieval (Danker and Anderson, 2010; Moscovitch et al., 2016) and shows age-related deficits in activity and connectivity during retrieval (Daselaar et al., 2006; Trelle et al., 2019; Deng et al., 2020). Thus, we hypothesized that *aging is associated with impaired mnemonic representations in early visual cortex and hippocampus but spared, or possibly even enhanced, mnemonic representations in the ATL* (**Hypothesis 2**).

We investigated Hypothesis 2 using a reactivation fMRI paradigm (Danker and Anderson, 2010; Rugg and Vilberg, 2013). As illustrated by **Fig. 1b**, during encoding scans, samples of YAs and OAs viewed 96 pictures of scenes paired with labels, and during retrieval scans, they recalled the scenes in response to the labels and rated the quality of their memories. These in-scan ratings were validated with a post-scan forced-choice memory recognition test, which showed that greater in-scan ratings were associated with better post-scan accuracy (**Fig. 1c**; see the Materials and Methods for more details on participants and the experimental design). Previous reactivation fMRI studies with OAs compared broad categories of stimuli (Abdulrahman et al., 2017; Johnson et al., 2015; Thakral et al., 2019; Wang et al., 2016) or presented stimuli multiple times during encoding (St-Laurent et al., 2014), precluding reactivation measures for individual events. In contrast, we measured the reactivation of individual events (each scene) by directly measuring *encoding-retrieval similarity* (ERS) in activation patterns (Ritchey et al., 2013; Wing et al., 2014). On the bases of Hypothesis 2, we predicted that, compared to YAs, OAs would show reduced ERS in the early visual cortex but spared (or even enhanced) ERS in the ATL.

## 2. Materials and Methods

### 2.1. Study Participants

Our study sample included 22 YAs and 22 OAs. One YA and one OA were excluded from analysis because of functional data missing from the first fMRI run due to a technical error. Another OA was excluded from analysis due to a poor quality T1 image, not allowing the participant’s functional images to be properly normalized into MNI space. This left a study sample of 21 YAs (12 women, age range = 18-30 years, *M* = 23.5 years, *SD* = 3.0 years) and 20 OAs (9 women, age range = 61-82 years, *M* = 70.5 years, *SD* = 5.4 years). Participants self-reported to be free of significant health problems (including atherosclerosis, neurological and psychiatric disorders), and not taking medications known to affect cognitive function or cerebral blood flow (except antihypertensive agents). Also, all participants were right-handed and completed at least 12 years of education. The OAs were additionally screened for dementia via the Mini-Mental State Examination (MMSE; inclusion criterion ≥ 27; M = 29.2, SD = 0.7; Folstein et al., 1975); no exclusions were necessary based upon this criterion. After study completion, participants were monetarily compensated for their time. Study results from the sample of YAs were previously reported in other manuscripts (Geib et al., 2017; Wing et al., 2014). The Duke University Institutional Review Board approved all experimental procedures, and participants provided informed consent prior to testing.

### 2.2. Experimental Design

Participants completed three encoding runs followed by three retrieval runs. During the encoding runs, participants explicitly studied a total of 96 color pictures of complex scenes (32 images per run, order randomized within run). The 96 images were very similar to the ones used to train the DNN model employed (VGG16), including many of the same images. During each encoding trial (4 sec), participants were presented a single picture with a unique descriptive label below the image (e.g., “tunnel” or “barn”). Within the trials, participants were asked to rate, on a four-point scale, the quality of the image (i.e., how well the image represents the label, 1 = *low quality*, 4 = *high quality*). This was to ensure participants would pay attention to the details of each image. Each encoding trial was followed by an active baseline interval of 8 sec, in which participants were presented digits from 1 to 4 and pushed the button corresponding to the presented numbers.

The retrieval runs were identical in format to the encoding runs, except the pictures of the scenes were not presented. During each retrieval trial, participants were presented the 96 descriptive scene labels previously presented along with the pictures of the scene, and participants were instructed to recall the corresponding image from encoding with as much detail as possible. Participants then rated, on a four-point scale, the amount of detail with which they could remember for the specific picture (1 = *least amount of detail*, 4 = *highly detailed memory*).

Immediately after the retrieval runs, participants completed a four-alternative forced-choice recognition test outside the scanner, in a testing room located adjacent to the MR scanner. For each recognition trial, the participants selected the picture they believed they saw during encoding among 4 pictures (1 target picture, 3 distractors) that were simultaneously presented for 5 sec (**Fig. 1b**, right). Participants then reported their confidence in the recognition decision using a 4-pt scale (1 = guess, 4 = very confident) in each trial. The performance of post-scan memory recognition for each participant was measured as the hit rate (the number of correct trials over the number of total trials).

### 2.3. MRI Data Acquisition

MRI data were collected on a General Electric 3T MR750 whole-body 60 cm bore MRI scanner and an 8-channel head coil. The MRI session started with a localizer scan, in which 3-plane (straight axial/coronal/sagittal) localizer faster spin echo (FSE) images were collected. Following, using a SENSE spiral-in sequence (repetition time [TR] = 2000 msec, echo time = 30 msec, field of view [FOV] = 24 cm, 34 oblique slices with voxel dimensions of 3.75 × 3.75 × 3.8 mm^3^), the functional images were acquired. The functional images were collected over six runs – three encoding runs and three retrieval runs; there was also a functional resting-state run after the third encoding run, which is not reported here. Stimuli were projected onto a mirror at the back of the scanner bore, and responses were recorded using a four-button fiber-optic response box (Current Designs, Philadelphia, PA, USA). Following, a high-resolution anatomical image (96 axial slices parallel to the AC-PC plane with voxel dimensions of 0.9 × 0.9 × 1.9 mm) was collected. Finally, diffusion-weighted images were collected, which are not reported here. Participants wore earplugs to reduce scanner noise, and foam pads were used to reduce head motion, and, when necessary, participants wore MRI-compatible lenses to correct vision.

### 2.4. Functional MRI Data Preprocessing

For each run, the first six functional images were discarded to allow for scanner equilibrium. All functional images were preprocessed in a SPM12 (London, United Kingdom; http://www.fil.ion.ucl.ac.uk/spm/) pipeline. Briefly, the functional images were slice timing corrected (reference slice = first slice), realigned to the first scan in the first session, and subsequently unwarped. Following, the functional images were coregistered to the skull-stripped high-resolution anatomical image (skull-stripped by segmenting the high-resolution anatomical image and only including the gray matter, white matter, and cerebrospinal fluid segments). The functional images were normalized into MNI space using DARTEL (Ashburner, 2007); the study specific high-resolution anatomical image was created using all of the study participants. The voxel size was maintained at 3.75 × 3.75 × 3.8 mm^3^ and the normalized-functional images were not spatially smoothed. Lastly, the DRIFTER toolbox (Sarkka et al., 2012) was used to denoise the functional images. We also calculated the temporal signal-to-noise ratio (SNR), that is, the ratio between mean and the standard deviation of the time series in a given ROI (Welvaert and Rosseel, 2013).

### 2.5. Functional MRI Analysis

#### 2.5.1. Functional Representational Similarity

To obtain the beta estimates for each event, we conducted a single-trial model analysis within a general linear model. These beta estimates were calculated using a least squares-separate approach (Mumford et al., 2012). This approach estimates a first-level model in which one regressor models a specific event of interest and another regressor models all the other events (each run included a regressor modeling these other trials). Each event was modeled with a stick function placed at stimulus onset convolved with a standard hemodynamic response function with the temporal and dispersion derivative. Each model also included the six raw motion regressions, a composite motion parameter (derived from the Artifact Detection Tools [ART]), outlier TRs (scan-to-scan motion > 2.0 mm or degrees, scan-to-scan global signal change > 9.0 *z* score; derived from ART), the white matter timeseries, and cerebrospinal fluid timeseries. In each model we also modeled the temporal and dispersion derivatives and implemented a 128 sec cutoff high-pass temporal filter.

These beta-images were used for (1) the representational similarity analysis combined with DNNs and (2) ERS. These analyses were conducted using in-house MATLAB (Natick, MA, USA) scripts (https://github.com/brg015). For the ERS analyses, we excluded trials in which participants responded (during retrieval) either not at all or in less than 250 msec.

#### 2.5.2. Representational Similarity Analysis Combined with Deep Convolutional Neural Networks

To examine our first goal, we performed representational similarity analysis (Kriegeskorte and Kievit, 2013; Kriegeskorte et al., 2008) based on stimuli models derived from DNNs (Khaligh-Razavi and Kriegeskorte, 2014; Kriegeskorte, 2015; Leeds et al., 2013; Wen et al., 2017) that captured the similarities between the stimuli in two different aspects (i.e., sensory vs. categorical aspects). The stimuli models (96 × 96 matrix) were correlated with the brain activation patterns similarity matrix (96 × 96 matrix) derived from searchlight volumes using Spearman’s correlation (a.k.a. model-brain fit), and the level of such model-brain fit may indicate the extent the neural representation reflects the processing of features associated with the given model (Kriegeskorte et al., 2006).

For the searchlight analysis across brain regions, a 5 × 5 × 5 voxel cube (Wing et al., 2014) was placed around a voxel location and the activation values from this cube were extracted and vectorized for each beta image, representing the local activation pattern associated with each stimuli at this voxel location (**Fig. 1a**). This procedure was conducted for each stimulus and the activation values from each stimulus were correlated (Fisher-transformed Pearson’s *r*) with each other, representing the brain activation patterns. The brain activation patterns were then correlated (Spearman’s correlation) with the DNN-stimuli models (model-brain fit), which was the value placed in the voxel location. This procedure was repeated for every voxel in the brain and the output of this analysis was searchlight volumes representing brain activation pattern-DNN layer stimuli model similarity.

For the stimuli model, we used a popular DNN known as VGG16 (Simonyan and Zisserman, 2014), which is pre-trained on approximately 1.8 million images of scenes in service of categorizing the images into 365 scene categories (Zhou et al., 2017). The VGG16 consists of 13 convolutional layers and 3 fully-connected layers. The first hidden DNN layer is a convolutional layer whose artificial neurons directly receive image inputs and have small spatial receptive fields. This layer is sensitive to low-level visual features, such as Gabor patches, boundaries, and blobs (Eickenberg et al., 2016). In comparison, the penultimate layer is the last hidden layer with 4096 artificial neurons, whose collective activation pattern is sent to the output layer (a.k.a. Softmax layer) to produce evidence scores of each of the 365 scene categories. Thus, the penultimate layer contains a rich array of computational information that is not defined by one visual feature (e.g., horizontal lines) or a simple combination of a few features (e.g., red horizontal lines) but instead a high-dimensional feature matrix which may dissociate many object classes (here, 365 classes), based on complex spatial configurations of information associated with different objects (ships, house, etc.) and backgrounds (sea, farmland, etc.) that allow to distinguish one scene from another (e.g., seaport vs. barn). In other words, the representation in the VGG16 penultimate layer directly supports its categorical outputs, which are highly consistent with the categorical judgements from human.

Therefore, we created stimuli models from the first hidden DNN layer (reflecting sensory features) and penultimate DNN layer (reflecting categorical features). These stimuli models were constructed by feeding the study stimuli through the pretrained DNN and for each stimulus at each layer of interest (i.e., the first and penultimate layers), extracting the activation values. For each layer of interest, the activation values between stimuli were correlated (Pearson correlation) with each other. This yielded two 96×96 matrices (one for each layer of interest), which represent the similarity of the DNN activation values and are the image models (sensory and categorical image models). It should be noted that, although the first convolutional layer mimics low-level visual processing, and the penultimate layer, categorical judgement, the whole VGG16 should not be viewed as a model that emulates the entire ventral visual pathway.

After conducting the searchlight analysis examining model-brain fit, the searchlight volumes were spatially smoothed with a 5 mm Gaussian kernel (Clarke et al., 2016; Clarke and Tyler, 2014). Then, for each model, we extracted model-brain fit from *a priori* ROIs derived from the AAL atlas (Tzourio-Mazoyer et al., 2002), which consisted of early visual cortex (bilateral calcarine, cuneus, and lingual ROIs) and ATL (left dorsal temporal pole and ventral temporal pole). We chose to examine only the left ATL because of our interest in categorical representations and an extensive literature demonstrating greater processing of categorical-related features (e.g., conceptual processing) in the left hemisphere (Hodges et al., 1992; Tyler et al., 2004; Warrington and McCarthy, 1983). See the Introduction for an explanation of *a priori* ROI choice.

#### 2.5.3. Encoding-Retrieval Similarity

To examine our second goal, we calculated ERS for each item using a searchlight procedure (Kriegeskorte et al., 2006). For each item (e.g., ‘barn’), its corresponding encoding and retrieval activation patterns were extracted from searchlight spheres, forming an encoding-retrieval vector pair. A 5 × 5 × 5 voxel cube was placed around each voxel and the activation patterns were extracted and vectorized. For each item, the encoding and retrieval vectors were correlated and the correlation value (Fisher-transformed Pearson’s *r*, which is ERS) was placed in the original center voxel location. Within each voxel location, the ERS value for each stimulus was averaged across all 96 encoding items. This procedure was repeated for every voxel within the brain. Afterwards, the searchlight volumes were spatially smoothed with a 5 mm Gaussian kernel (Clarke et al., 2016; Clarke and Tyler, 2014). We then extracted ERS from *a priori* ROIs derived from the AAL atlas (Tzourio-Mazoyer et al., 2002), which consisted of early visual cortex (bilateral calcarine, cuneus, and lingual ROIs), ATL (left dorsal temporal pole and ventral temporal pole), and the hippocampus (bilateral hippocampi). See the Introduction for an explanation of *a priori* ROI choice. In addition to item-level ERS, as a control analysis, we also calculated set-level ERS, which is ERS between an item and every other item within the stimuli set (Ritchey et al., 2013; Wing et al., 2014), reflecting the baseline level of recurring activation patterns that were non-specific to the retrieval of specific item. Thus, the value of dissociating item- and set-level ERS is that we are able to control for the amount of pattern similarity that would be seen between any pair of items, and describe a more specific reinstantiation of a neural pattern unique to the target item. The analysis contrasting item and set ERS was constrained to the items that were subsequently remembered on the post-scan memory task.

### 2.6. VGG16 Model Validation

We conducted two preliminary analyses to validate the VGG16 (Simonyan and Zisserman, 2014) as a model of sensory to categorical representations in our study. (i) The VGG16 was already successfully trained to classify 1.8 million scenes into 365 categories (Zhou et al., 2017), but we wanted to confirm it could also classify the 96 images employed in our study. Given that some of the scene labels we used were different than the categories used to train the VGG16 but were nevertheless closely related terms (e.g., “seaport” in Fig. 1-B, versus “harbor” in the VGG16 output category), it is difficult to evaluate the performance of VGG16 directly. Therefore, we created a slightly modified VGG16 for binary indoor-outdoor scene classification. This was achieved by removing the last layer of the VGG16 and adding a layer with two outputs (with a Softmax activation function), corresponding to indoor and outdoor scenes. The revised VGG16 was then trained using three pictures from each image category (images from the post-scan recognition task besides the target images) and tested on the pictures presented within the scanner. After 30 epochs, the revised VGG16 was able to classify the images into indoor vs. outdoor domains with 94.8% accuracy, suggesting that the stimuli images were well-matched with the training images. It should be noted that the model with this binary indoor-outdoor scene classification was only used to validate the use of the VGG16 within our study; the original pretrained VGG16 was used for stimuli model construction. (ii) We correlated the sensory and categorical stimuli models based on the original pretrained VGG16 and found that they were only moderately correlated with each other (*r* = 0.32, Figure S1), indicating that each model represents unique features. In comparison, the stimuli models constructed from adjacent layers are more closely correlated with each other (**Supplementary Fig. 1**).

### 2.7. Statistical Analysis

Statistical analyses (unless otherwise stated) were conducted within StatsModels (Seabold and Perktold, 2010) ran in Python 3 (Python Software Foundation, https://www.python.org/). Repeated measures ANOVAs were used to assess omnibus effects. Two-sided linear mixed effects models, with participant specified as a random effect, were used as a substitute of t-test or correlation. This model allowed us to include SNR as a nuisance variable when needed, as ATL can be vulnerable to low SNR. The significance of effects was based on the z-scores associated with the parameter estimates. Before entering the data into the linear mixed effects models, all values were *z*-transformed. All effect sizes reported in the manuscript were Cohen’s *d*.

## 3. Results

### 3.1. Behavioral Results

We started by investigating the memory performance of the two age groups. In the post-scan recognition test, OAs showed lower number of hits comparing with YAs (YA: 75.10±10.98, OA: 61.00±8.26, t=4.63, p=4.02e-5), as well as lower recognition confidence (YA: 3.11±0.30, OA: 2.74±0.41, t=3.31, p=0.0022), suggesting age-related memory decline. In the retrieval scan, the two age groups showed comparable level of self-report recall vividness inside scanner (YA: 2.72±0.46, OA: 2.70±0.40, t=0.12, p=0.91). The post-scan memory accuracy (hit rate) was positively related to vividness rating during retrieval **(Fig. 1c)**, as is indicated by a significant effect of vividness (F(3,117)=13.17, p<2e-7). In addition, the confidence level of post-scan recognition was also significantly contributed by in-scan vividness (**Fig. S2**; F(3,117)=64.62, p=2e-16). Memory performance for each stimulus can be found in **Supplementary Table 2**.

### 3.2. Perceptual Representations: DNN-based Representational Similarity Analysis

Our first hypothesis was that age-related visual dedifferentiation impairs the differentiation of sensory features in early visual cortex but not categorical features in the ATL, which may even be enhanced in aging. Using DNN-based representational similarity analysis, we tested this hypothesis by comparing age-related differences in model-brain fit for the sensory model based on the first VGG16 layer, and for the categorical model based on the penultimate VGG16 layer. We tested this hypothesis in two *a priori* ROIs – early visual cortex and ATL. For raw model-brain fit values (see **Supplementary Fig. 3a**), we performed an omnibus *Model_Type* (sensory/categorical) × *ROI* (early visual cortex/ATL) × *Age* (YA/OA) repeated measures ANOVA, and we found a significant main effect of *ROI* (F(1,39)=15.37, p<0.001) and a significant *Age × ROI* interaction (F(1,39)=12.14, p<0.005), while the main effect of *Model_Type* (F(1,39)=0.34, p=0.57), *Model_Type × Age* interaction (F(1,39)=1.66, p=0.21), *Model_Type × ROI* interaction (F(1,39)=0.095, p=0.76) and *Model_Type × ROI × Age* interaction were not significant (F(1,39)=1.82, p=0.185). In an ROI-level analysis, as illustrated in **Fig. 2a**, which shows *z*-scored 2^nd^ order correlations, the evidence was consistent with our first hypothesis: compared to the YAs, in early visual cortex, the sensory model-brain fit was reduced in the OAs (*β* = −0.39, *z* = 19.86, *p* < .0001, *d* = 0.83), whereas in the ATL, the categorical model-brain fit was enhanced in the OAs (*β* = 0.26, *z* = 12.82, *p* < .0001, *d* = 0.53). In other words, whereas early visual cortex showed age-related *de*differentiation, the ATL showed age-related *hyper*differentiation. As the ATL is particularly vulnerable to low signal-to-noise ratio (SNR), we repeated the analysis statistically controlling for SNR and still found that in the ATL that categorical model-brain fit was enhanced in the OAs compared to YAs (*β* = 0.26, *z* = 15.61, *p* < .0001). We did not find statistically significant age-group differences of the sensory model-brain fit in the ATL (*β* = 0.17, *z* = 1.39, *p* = .17, *d* = 0.33), but we did find, compared to YAs, that the categorical model-brain fit in the early visual cortex was reduced in the OAs (*β* = −0.27, *z* = 3.25, *p* < .01, *d* = 0.54). In sum, this is, to our knowledge, the first evidence of age-related hyperdifferentiation of activation patterns in the ventral pathway or any brain region.

**Figure 2:**
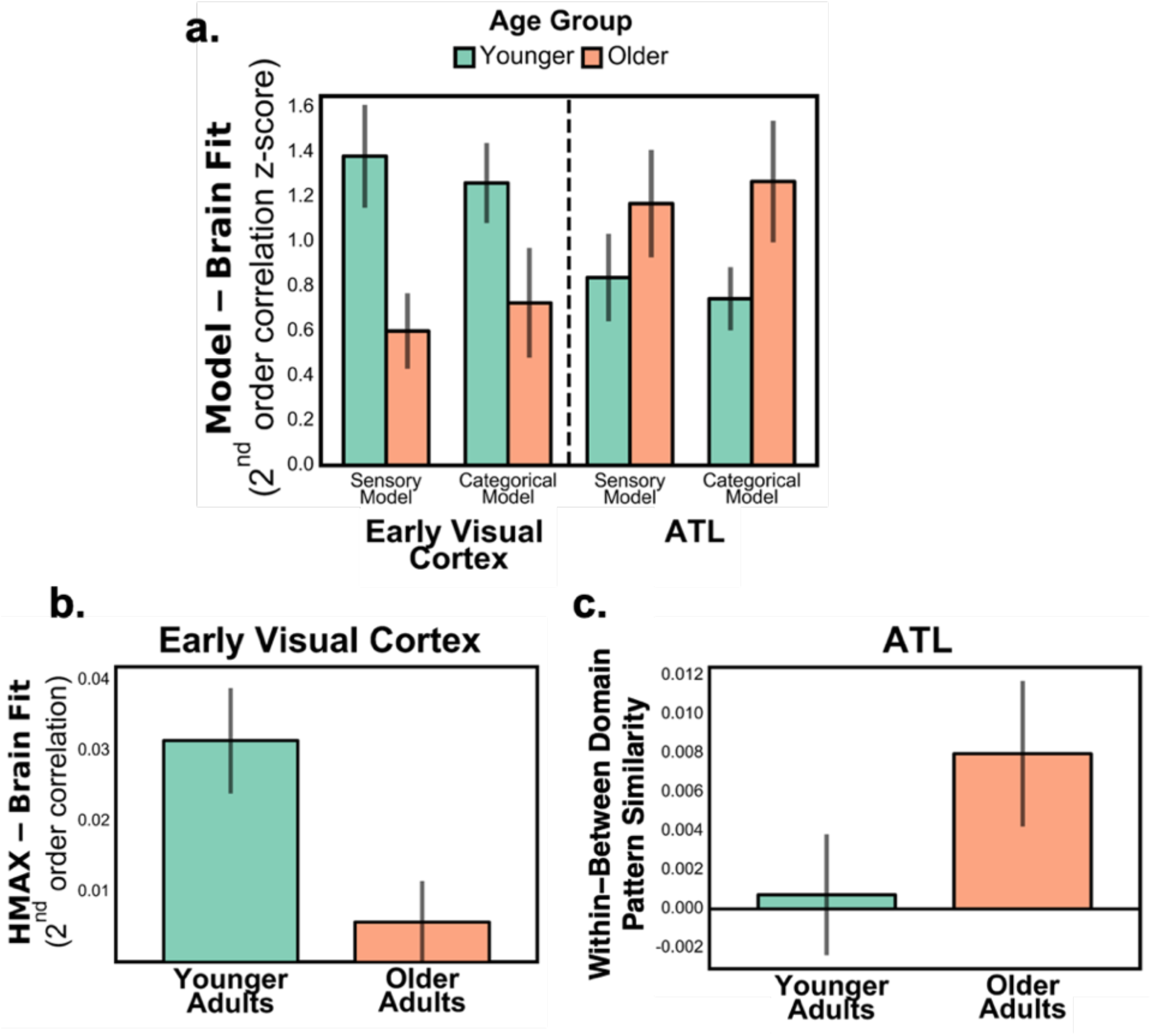
Age-related neural dedifferentiation and hyperdifferentiation. **Panel a** shows within the early visual cortex and ATL the stimuli model (sensory and categorical)-brain fit; the figure shows z-scored (mean set to one) 2^nd^ order correlations. Compared to the YAs, in the early visual cortex, we found that the sensory model-brain fit was reduced in the OAs (age-related dedifferentiation), whereas in the ATL, the categorical model-brain fit was enhanced in the OAs (age-related hyperdifferentiation). **Panel b** shows the HMAX C1 response model-brain fit in early visual cortex. We found that, compared to YAs, OAs exhibited reduced HMAX-brain fit in early visual cortex. **Panel c** shows the brain activation pattern similarity for within-domain minus between-domains (domains = indoor vs. outdoor scenes) in the ATL. We found that, compared to the YAs, OAs exhibited enhanced activation pattern similarity for within-than between-domains in the ATL. Error bars represent the standard error of the mean. Error bars represent the standard error of the mean. ATL = anterior temporal lobe, OAs = older adults, YAs = younger adults.

Although DNNs provide stronger models of visual representations in the ventral visual pathway than traditional theoretical models (e.g., HMAX, object-based models; Cadieu et al., 2014; Groen et al., 2018), DNNs are sometimes critiqued for being too complex and, therefore, less interpretable. We believe that this level of complexity is necessary to map representations within the brain (Kriegeskorte and Douglas, 2018), but the critique of interpretability is well received. Therefore, we examined if the age-related differences can also be observed using more traditional models. (i) For sensory representations, we used the C1 responses of the HMAX model (Clarke and Tyler, 2014; Serre et al., 2007), which was proposed to reflect properties of early visual cortex (Riesenhuber and Poggio, 1999; Serre et al., 2007). Consistent with the DNN analysis, compared to the YAs, the OAs exhibited reduced HMAX-based model-brain fit in early visual cortex (**Fig. 2b**; *β* = −0.39, *z* = 13.73, *p* < .0001, *d* = 0.82). In addition, the HMAX-based model-brain fit was strongly correlated with the early DNN-based model-brain fit across participants (r = 0.74, p < 0.001), suggesting good consistency of the two measurements. (ii) For categorical representations, we examined brain activation pattern similarity for within-compared to between-domains (indoor vs. outdoor scene trials), since images within the same domain likely share large amounts of objects and visual features, as demonstrated in our preliminary analysis using the VGG16 to classify indoor and outdoor scenes (see Preliminary analyses in the Results). We performed a *Model_Type* (HMAX C1/within-minus-between domains) × *ROI* (early visual cortex/ATL) × *Age* (YA/OA) ANOVA on these two alternative model-brain fit measures. We found significant main effect of *ROI* (F(1,39) = 5.23, p < 0.05), *Age* × *ROI* interaction (F(1,39) = 9.61, p < 0.005), and *Model_Type* (F(1,39) = 4.89, p < 0.05). The *Model_Type* × *Age* interaction (F(1,39) = 3.51, p < 0.1), *Model_Type* × *ROI* interaction (F(1,39) = 3.52, p < 0.1) and *Model_Type* × *ROI* × *Age* interaction (F(1,39) = 3.52, p < 0.1) were non-significant, although noticeable to some extent. Consistent with the DNN analysis, compared to the YAs, the OAs exhibited enhanced activation pattern similarity for within-than between-domains in the ATL (**Fig. 2c**; *β* = 0.23, *z* = 3.36, *p* < .001, *d* = 0.46; see **Supplementary Fig. 3b** for pattern similarity values within- and between-domains). These findings supported the robustness of DNN-based analysis on perceptual representation, providing further evidence for age-related dedifferentiation of sensory features in early visual cortex and hyperdifferentiation of categorical features in ATL.

As an exploratory analysis, we further investigated the links between encoding representations and memory (**Supplementary Table 3**). In YAs, we found a significant positive correlation between post-scan memory accuracy and sensory model-brain fits in early visual cortex (DNN Layer 1 model-brain fit: r = 0.45, p < 0.05; HMAX C1 model-brain fit: r = 0.51, p < 0.05), suggesting the sensory representation in visual cortex supported later memory retrieval. In comparison, in OAs, such a correlation between memory and sensory model-brain fits was absent (DNN Layer 1 model-brain fit: r = 0.04, p > 0.5; HMAX C1 model-brain fit: r = −0.03, p > 0.5), although there were no significant differences between the correlations of the two age groups. Collapsing across all participants (YAs & OAs), the association between early sensory model-brain fits for early layer models (DNN Layer 1, HMAX C1) and memory accuracy was significant (DNN Layer 1 model-brain fit: r = 0.46, p < 0.005; HMAX C1 model-brain fit: r = 0.47, p < 0.005); when corrected for age effect, such behavioral correlate was still significant for the HMAX C1 model (partial correlation, r=0.32, p=0.042) and almost reaching significant for the DNN Layer 1 model (partial correlation, r=0.31, p=0.051). We did not find significant correlation between memory and categorical model-brain fits in ATL in either YAs (r=0.14, p=0.54) or OAs (r=0.053, p=0.83).

Although the *a priori* ROI analysis was used to test our hypothesis, as an exploratory analysis, we conducted the whole-brain searchlight analysis. Within both age groups, the first layer of the VGG16 was primarily associated with early visual regions, and the penultimate layer was additionally linked to anterior temporal, parietal, and frontal regions (**Fig. 3**; see **Supplementary Table 1** for cluster coordinates). This is consistent with previous research that early DNN layers identified posterior brain regions mediating sensory representations, and later layers, anterior brain regions mediating categorical representations (Güçlü and van Gerven, 2015; Khaligh-Razavi and Kriegeskorte, 2014; Kriegeskorte, 2015; Leeds et al., 2013; Wen et al., 2017). With respect to age-related differences, YAs showed stronger sensory model-brain fit in the early visual regions, whereas OAs showed stronger categorical model-brain fit in anterior parahippocampal gyrus and some frontal regions (**Supplementary Table 1**). In sum, the whole-brain searchlight analysis largely mirrors the *a priori* ROI analysis.

**Figure 3:**
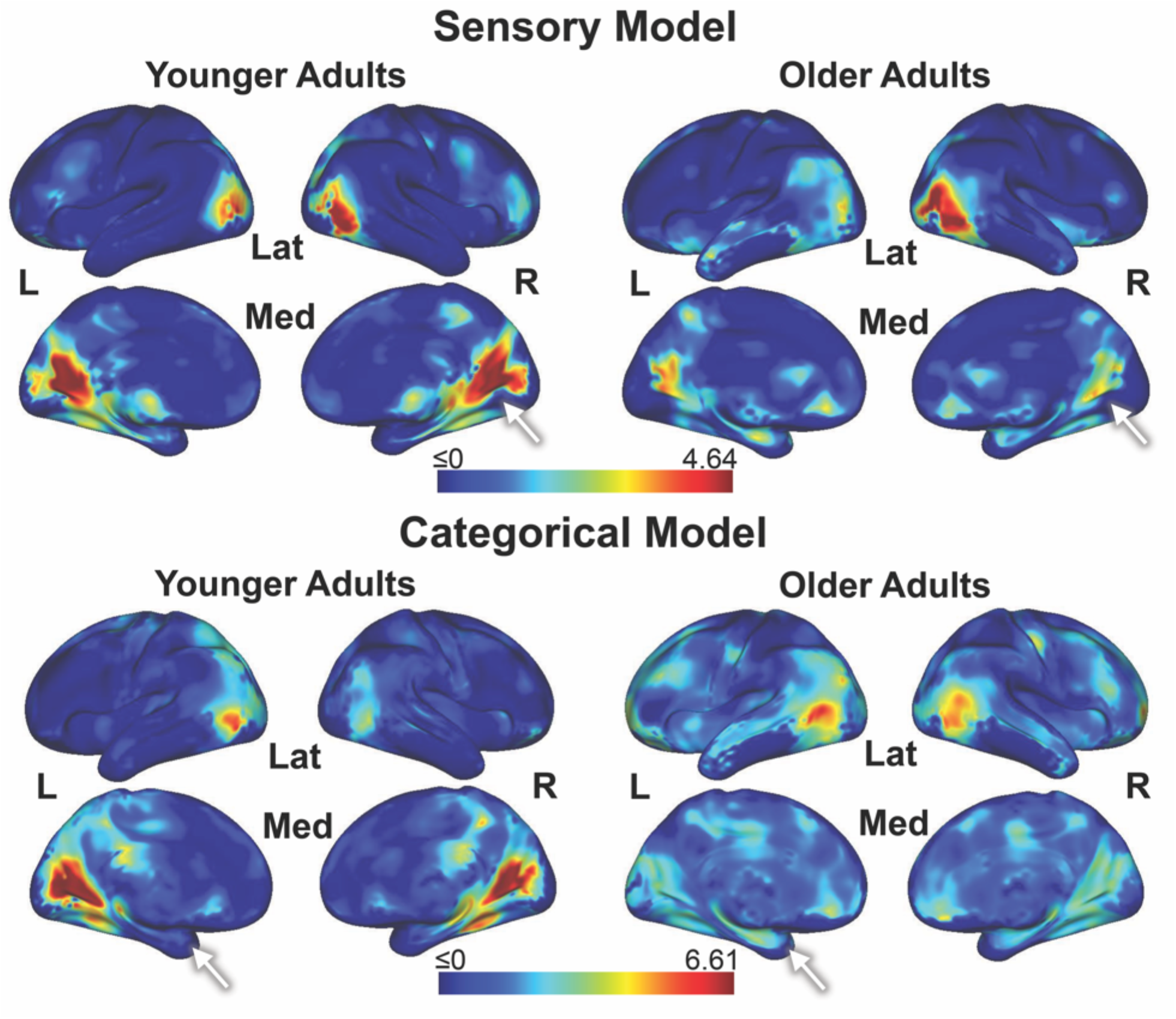
Sensory and categorical model-brain fit. The figure shows the whole-brain searchlight analysis of the stimuli model (sensory and categorical)-brain fit. Within both the YAs and OAs, we found that the sensory model (based upon the first DNN layer) correlated predominately with brain activation patterns in the visual cortices and the categorical model (based upon the penultimate DNN layer) additionally correlated with brain activation patterns in anterior temporal, parietal, and frontal regions. Qualitatively, in these maps it can be seen that that the sensory model is more strongly associated with earlier visual cortex region activation patterns in YAs than OAs, whereas the categorical model is more strongly associated with more anterior ventral visual pathway region, such as the ATL, activation patterns in OAs than YAs (see white arrows). DNN = deep convolutional neural network; L = left; Lat = lateral; Med = medial; OAs = older adults; R = right; YAs = younger adults.

### 3.3. Mnemonic Representations: Encoding-Retrieval Similarity

Our second hypothesis was that aging impairs mnemonic representations for sensory features in early visual cortex and hippocampus but not for categorical features in the ATL. To test this hypothesis, we calculated encoding-retrieval similarity (ERS; Ritchey et al., 2013; Wing et al., 2014). We tested this hypothesis using the same *a priori* ROIs used to test our first hypothesis with the addition of the hippocampus because of the second hypothesis’s relation to memory (see the Methods for more details). Consistent with Hypothesis 2, as shown within **Fig. 4a**, we found that, compared to YAs, OAs exhibited reduced ERS in the early visual cortex (*β* = −0.35, *z* = 14.34, *p* < .0001, *d* = 0.74) and hippocampus (*β* = −0.45, *z* = 3.37, *p* < .001, *d* = 0.97), but increased ERS in the ATL (*β* = 0.20, *z* = 3.62, *p* < .001, *d* = 0.39; see **Supplementary Fig. 3c** for raw ERS values). As the ATL is particularly vulnerable to low SNR, we repeated the analysis statistically controlling for SNR and still found the same effect within the ATL (*β* = 0.24, *z* = 2.98, *p* < .01). In sum, these results indicated age-related decrease in mnemonic representations within regions associated with sensory features, as well as age-related increase in mnemonic representations within regions associated with categorical features.

**Figure 4:**
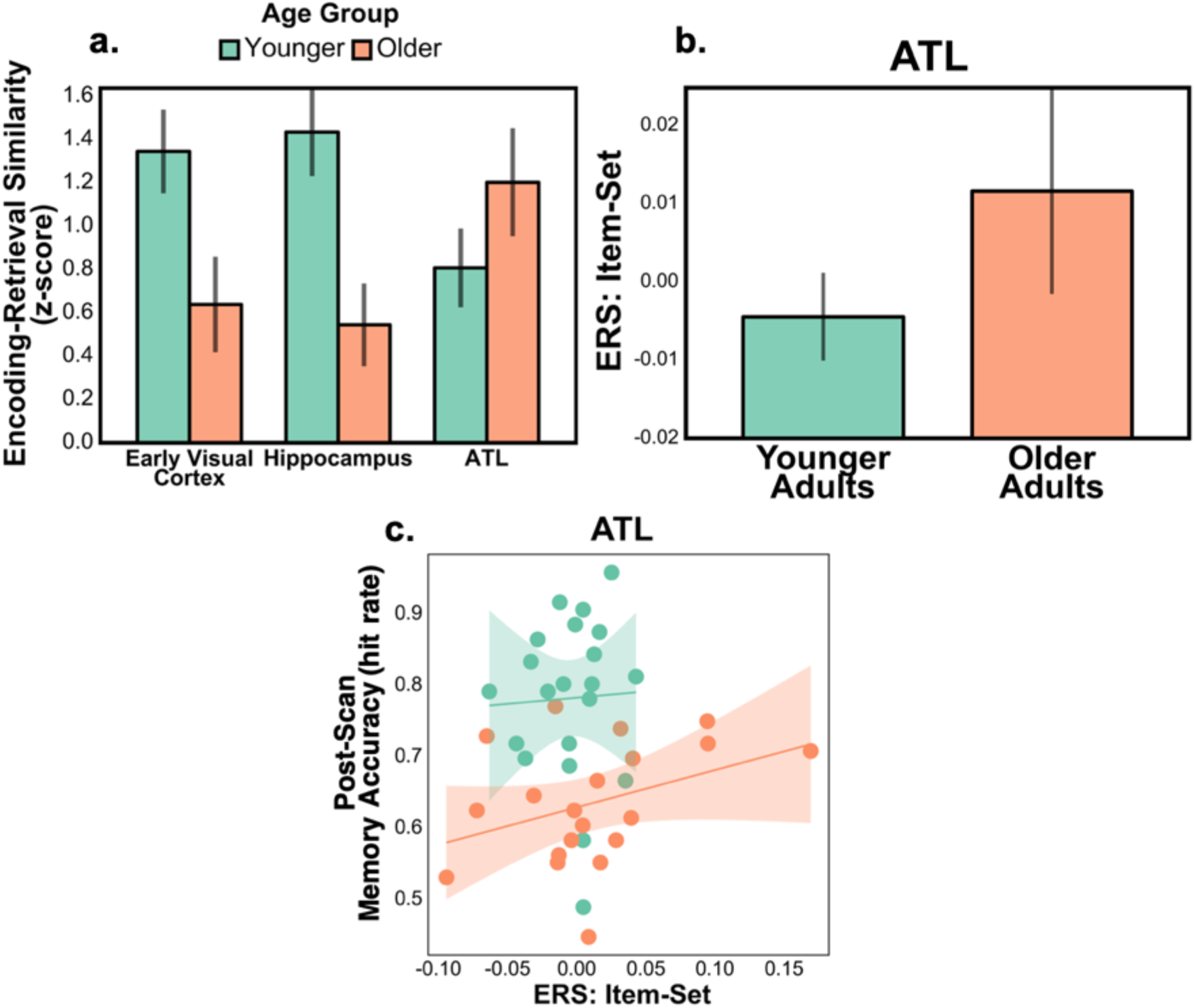
Age-related differences in mnemonic representations. **Panel a** shows age-related differences in ERS. We found, compared to the YAs, in the early visual cortex and hippocampus that the OAs exhibited reduced ERS, whereas in the ATL, the OAs exhibited increased ERS. The figure shows z-scored (mean set to one) ERS; error bars represent the standard error of the mean. **Panel b** shows item-specific ERS (item-minus-set level ERS) in the ATL for trials that were subsequently remembered on the post-scan memory recognition task. We found that, compared to the YAs, OAs exhibited enhanced item-specific ERS in the ATL. Error bars represent the standard error of the mean. **Panel c** shows the relation between item-specific ERS in the ATL (limited to the trials that were subsequently remembered) and accuracy of the post-scan memory recognition task. We found in the OAs that increased item-specific ERS in the ATL was associated with better accuracy of the post-scan memory recognition task; translucent bars around the regression lines represent the 95% confidence intervals. ATL = anterior temporal lobe; ERS = encoding-retrieval similarity; OAs = older adults; YAs = younger adults.

In order to determine whether the dedifferentiation of sensory representations observed in early visual cortex were associated with a deficit in mnemonic representations, we examined the correlation between the sensory-model brain fit during encoding and ERS in early visual cortex. We found a significant positive correlation (all participants, age-controlled partial correlation: r = 0.39, p = 0.01; YA: r = 0.36, p = 0.11; OA: r = 0.46, p = 0.042). This finding supports the assumption that age-related perceptual dedifferentiation is associated with a deficit in mnemonic representations.

Within the previous analysis we examined ERS for individual items (i.e., the similarity between pairs of encoding and retrieval activation patterns of the same scenes; item-level ERS). However, in order to make the claim that such retrieval reactivation is related to the specific memory items during encoding, it is necessary to also compare item-level ERS with set-level ERS (i.e., average encoding-retrieval pattern similarity between one scene and all other scenes in the stimuli set, which reflects baseline neural reactivation; Koen and Rugg, 2016; Ritchey et al., 2013; Wing et al., 2014). Furthermore, in order to strengthen our claim that ERS here reflects memory, it is necessary to repeat the analysis only on items that were subsequently remembered on the post-scan memory recognition task. Therefore, we repeated the analysis subtracting set-level ERS from item-level ERS and only including trials that were subsequently remembered on the post-scan memory task. The only region that still exhibited the same age group difference pattern was the ATL, in which, compared to YAs, OAs exhibited enhanced ERS in the ATL (**Fig. 4b**; *β* = 0.016, *z* = 2.59, *p* = .01, *d* = 0.37; see **Supplementary Fig. 3d** for raw ERS values). The age-related increase in item-specific ERS in the ATL suggests that this region contributes to memory to a greater extent in OAs than YAs. To investigate this idea, we examined the relation between item-specific ERS (item-minus-set ERS) in the ATL and performance on the post-scan memory recognition task. As illustrated in **Fig. 4c**, greater ATL item-specific ERS was associated with better accuracy of the post-scan memory recognition task in the OAs (*β* = 0.51, *z* = 2.17, *p* < .05) but not YAs (*β* = 0.082, *z* = 0.092, *p* = .93); it should be noted, however, that the age group by ERS interaction was not statistically significant (*β* = 0.40, *z* = 0.42, *p* = .68). As a control analysis, the item-specific ERS of subsequently forgotten items was not associated with better post-scan recognition accuracy in YAs (*β* = −0.10, *z* = −1.55, *p* = .12) or OAs (*β* = 0.44, *z* = 1.34, *p* =.18). Meanwhile, since such association with behavior was across participant, we further performed an Age (YA/OA) × Memory (Hits/Misses) repeated measures ANOVA on item-specific ERS in ATL. We did not find a higher ERS associated with hit items (F(1,39) = 0.02, p = 0.90) or a significant Age × Memory interaction (F(1,39) = 1.74, p = 0.19).

In sum, consistent with our second hypothesis, we found that OAs exhibited reduced ERS in the visual cortex and hippocampus but increased ERS in the ATL. The latter effect was found to be item-specific and related to better memory accuracy across OA participants.

## 4. Discussion

The overarching goal of the study was to examine age-related differences in the quality of perceptual and mnemonic representations. We had two main findings. First, in early visual cortex, activation patterns associated with sensory features showed dedifferentiation in OAs, replicating previous age-related dedifferentiation findings, whereas in the ATL, activation patterns associated with categorical features showed *hyper*differentiation in OAs. This is, to our knowledge, the first report of age-related hyperdifferentiation. Second, for mnemonic representations, we found that increased age was associated with impaired mnemonic representations (i.e., decreased ERS) in the early visual cortex and hippocampus but enhanced mnemonic representations (i.e, increased ERS) in the ATL. The enhanced mnemonic representations in the ATL was associated with better memory in OAs. These findings are discussed in greater detail below.

### 4.1. Age-related Neural Dedifferentiation

Previous studies have brought accumulating evidence for age-related neural dedifferentiation (Carp et al., 2011; Chee et al., 2006; Goh et al., 2010; Payer et al., 2006; Voss et al., 2008; Zheng et al., 2018; Sommer et al., 2019; Trelle et al., 2019; Bowman et al., 2019; Koen et al., 2019; Koen and Rugg, 2019; Koen et al., 2020). Age-related dedifferentiation has been often operationalized as reduced specificity of neural responses towards different stimuli sets (e.g., faces vs. places, living vs. non-living), which are heterogeneous domains with considerable within-domain variation. The current approach expands the utility of the research concept of dedifferentiation by considering how specific aspects of visual perception representations may be impaired in the visual system in older adults. Here, we compared age-related differences in the cortical differentiation of sensory vs. categorical features using a novel DNN-based representational similarity analysis approach (Güçlü and van Gerven, 2015; Khaligh-Razavi and Kriegeskorte, 2014; Kriegeskorte, 2015; Leeds et al., 2013; Wen et al., 2017). Consistent with our hypothesis, within early visual cortex, we found evidence for age-related neural dedifferentiation of sensory features, where, compared to the YAs, brain activation patterns in early visual cortex exhibited a worse fit with the sensory model in the OAs (Figs. 2a, b). It is well known that increased age is associated with a robust decline in visual performance (Monge and Madden, 2016; Owsley, 2011). Our finding suggests that this decline may reflect not only a decline in perceptual operations (*processes*) performed on sensory information coming from earlier regions in the ventral stream, but also a deficit in the quality of visual information itself (*representations*) (Carp et al., 2011).

The dedifferentiation observed in early visual cortex may have downstream influences on subsequent encoding-retrieval pattern matching, as indicated by the significant positive correlation between sensory model-brain fit and ERS within early visual cortex (all participants, age-controlled partial correlation: r = 0.39, p = 0.01; YA: r = 0.36, p = 0.11; OA: r = 0.46, p = 0.042). No correlation between sensory model-brain fit in early visual cortex and categorical model brain fit in ATL was found (all participants, age-controlled partial correlation: r = 0.017, p = 0.92; YA: r = −0.15, p = 0.52; OA: r = 0.15, p = 0.53), suggesting that the downstream hyperdifferentiation was not driven by dedifferentiation in visual cortex. Taken together, these results provide evidence that the quality of perceptual representations at encoding will affect later mnemonic reinstatement. We used a straightforward recognition test to evaluate for successful retrieval, but future studies may wish to examine what influence this representational shift has for age-related changes in strategic retrieval operations mediated by more anterior PFC networks. Although the current analysis did not include multivariate measures of connectivity between regions, such an analysis would provide for a fruitful means of determining how OA networks adapt to declining differentiation of neural signals in primary visual regions.

### 4.2. Age-related Neural Hyperdifferentiation

In contrast to sensory features in early visual cortex, we found that categorical features in the ATL were not just spared but actually enhanced within OAs. Within the ATL, compared to YAs, activation patterns had a better fit with the categorical model in the OAs (Fig. 2a). Thus, in contrast to age-related dedifferentiation in early visual cortex, in the ATL, we found age-related hyperdifferentiation. Regarding the regions that exhibited age-related hyperdifferentiation, the ATL is hypothesized to store prior knowledge (Lambon Ralph et al., 2017; Zhao et al., 2017). Given that OAs have a stronger knowledge network than YAs (Long and Shaw, 2000; Park et al., 2002), likely from more years of knowledge accruement, it is possible that certain processes, such as object recognition, are more reliant on categorical features. Perhaps the enhanced categorical model-brain fit in the ATL within the OAs reflects OAs utilizing enhanced semantic knowledge in service of perception.

Given that we operationalized dedifferentiation using RSA, as a reduction in the fit between activity patterns and a model of the stimuli (visual or categorical), hyperdifferentiation was conversely operationalized as an increase in this fit. The neural mechanisms of dedifferentiation are unclear, and hence by necessity, those of hyperdifferentiation are also uncertain. We believe that the hyperdifferentiation finding reflects age-related differences in the strategies used during encoding. Representations are not necessarily ‘hard-wired’ into the brain and the representational space can be warped in response to task demands (Çukur et al., 2013; Martin et al., 2018; Wang et al., 2018). During encoding, participants rate the representativeness of its picture for its label; we speculate that YAs paid more attention to visual features (e.g., *dark* forest, *shiny* bathroom, *tall* cathedral), whereas OAs paid more attention to scene-level conceptual features (e.g., *jungle-like* forest, *master* bathroom, *Gothic* cathedral). The latter strategy is consistent with a greater age-related reliance on gist-level representations of scenes (Gutchess et al., 2006), and would tend to enhance the fit of activation patterns with late DNN layers (i.e., close to the scene label), which is what we mean by hyperdifferentiation of categorical representation in the current study.

Furthermore, this age-related shift in encoding strategies is likely to emerge slowly over the lifespan; we found no correlation between the strength of corresponding sensory—early visual cortex and categorical—ATL representations (r = −0.08, p = 0.59), suggesting that it may not be feasible to observe this shift in a cross-sectional cohort. In order to confirm our interpretation of the results, a study manipulating participants’ use of sensory and categorical representations is necessary. It would be also important to investigate the phenomenon of hyperdifferentiation of conceptual information using models of stimulus semantics based on validated feature norms, rather than inferring category selectivity via DNNs. For example, in a recent RSA study (Davis et al., 2020) using object pictures instead of scenes, we decomposed semantic features into Observed (“is round”), Taxonomic (“is a fruit”), and Encyclopedic (“is sweet”) sub-categories based on an independently collected set of conceptual feature norms and a validated organization of feature categories (McRae et al., 2005). This type of analyses could build on the current DNN-based results by providing more information about the nature of the conceptual representations underlying the current age-related hyperdifferentiation finding, revealing perhaps the type of semantic information that is most relevant reliable memory formation in younger or older adults.

### 4.3. Age-related Differences in Mnemonic Representations

Our second question examined age-related differences in mnemonic representations. To examine this question, we examined the *reactivation* of mnemonic representations, which was assessed as the similarity between activation patterns during encoding and retrieval (ERS). Consistent with previous studies (Johnson et al., 2015; St-Laurent et al., 2014), compared to YAs, in OAs ERS was weaker in the early visual cortex (Fig. 4a). Given that we also found age-related dedifferentiation of sensory representations in this region, the age-related attenuation of ERS in early visual cortex likely reflects a negative impact of degraded sensory features on visual memories. It should be noted that even though there are studies that have also demonstrated age-related reductions in reactivation within visual cortex (Johnson et al., 2015; St-Laurent et al., 2014), some previous studies did not find this pattern (Abdulrahman et al., 2017; Thakral et al., 2019; Wang et al., 2016). The latter used multi-voxel pattern analysis (MVPA) to identify the reactivation of *classes of stimuli* (e.g., words vs. objects), whereas we used ERS to detect the reactivation of individual items of the same class. Thus, it is possible that MVPA is less sensitive than ERS in detecting age-related reactivation deficits. We also found an age-related ERS reduction in the hippocampus (Fig. 4a). This finding is consistent with evidence of impaired hippocampal activity in OAs (Kennedy et al., 2017; Nyberg, 2017), and it extends this evidence to multivariate activation patterns. Besides normal aging, the use of ERS to investigate hippocampal memory representations could be useful for investigating Alzheimer’s disease, in which the hippocampus structure and function are known to be particularly compromised (Frisoni et al., 2010; Dickerson and Sperling, 2008; Wang et al., 2006; Rathore et al., 2017).

Finally, we found that, compared to YAs, OAs exhibited enhanced ERS in the ATL (Fig. 4a). As the ATL has been associated with abstract semantic representations (Lambon Ralph et al., 2017; Zhao et al., 2017), enhanced ATL ERS in OAs may reflect a greater reliance on semantic knowledge in service of memory. This idea is consistent with our finding of age-related hyperdifferentiation for categorical features in this region (Fig. 2a). The accumulation of knowledge during the lifespan is likely to lead to more distinct and detailed semantic representations, which could partially counteract the degraded quality of sensory information flowing through the visual system (Monge and Madden, 2016).

### 4.4. Limitations and Further Considerations

Although the models chosen in this study were intended to measure sensory and categorical representations, we noted that there was an absence of reliable age-invariant model specificity, as suggested by an insignificant Model_Type × ROI interaction for the DNN-based results. There may be several explanations to this observation. Firstly, the DNN may not fully characterize the region-specific processing of perceptual information, which may be addressed by using improved models in future studies. Secondly, the sensory and categorical aspects of the stimuli may not be completely independent. Note that, the DNN-based sensory and the categorical models were moderately correlated (see 3.1), and it is common that scenes within a category may share similar low-level visual features (e.g., city skyline scenes may all have horizontal and vertical straight lines and sharp angles; rollercoaster scenes may all contain smooth spiral curvatures). In comparison, the analyses on alternative models (HMAX C1 vs. indoor-outdoor) showed improved Model_Type × ROI effect, probably due to the fact that scenes within the same broad domain (indoor or outdoor) were more heterogeneous in their low-level visual features, leading to better model specificity in different brain regions. Despite these limitations, our study points to the application of these computational models in quantifying sensory and categorical information, which could help future studies better control for the dependency between different feature similarities across stimuli items before carrying out the experiments.

We found that in OAs, increased item-specific ERS in the ATL was associated with better accuracy of the post-scan memory recognition task (Fig. 4c), supporting the importance of ATL representational quality for memory in aging. This across-subject correlation suggests that individual differences in memory performance may be at least partially mediated by neural representational quality. However, it is important to note that such association with behavior was across participant, whereas, within individuals, we did not find a higher ERS associated with hit items (F(1,39) = 0.02, p = 0.90) or a significant Age × Memory interaction (F(1,39) = 1.74, p = 0.19). Therefore, based on current evidence, caution is needed to interpret enhanced ERS in ATL as a compensatory mechanism in OAs. In addition, the post-scan memory test is a largely visual memory task that may not rely on the type of semantic processing we show to be centered in the left ATL. Therefore, future research may further elucidate the functional consequences of such age-related shift to ATL engagement by adopting a more diverse array of post-scan memory tests tapping both perceptual and conceptual memory.

### 4.5. Conclusions

The traditional idea that aging is associated with a generalized decline in cognitive abilities and their underlying neural mechanisms has been challenged by evidence that although some cognitive and brain mechanisms are impaired by aging, others are not only spared but even enhanced by aging (Long and Shaw, 2000; Park et al., 2002). One aspect of cognition that is spared by aging are processes that rely on categorical features, such as conceptual and semantic processing, but the neural mechanisms of these spared functions are largely unknown. The results of the present study suggest that one factor contributing to the preservation of these processes in old age is the enhanced quality of categorical representations in the ATL. These spared categorical representations may contribute not only to perceptual but also to mnemonic aspects of cognition. Lastly, these findings have implications not only for understanding normal aging but also the effects of pathological aging (e.g., Alzheimer’s disease) on memory representations, which, to our knowledge, has yet to be examined.

## 5. Acknowledgements

We thank four anonymous reviewers for their valuable comments. This work was supported by the National Institute on Aging (R01 AG019731 awarded to RC and F31 AG060691 awarded to ZAM) and National Institute of Mental Health (F31 MH114454 to BRG). The funding agency had no role in the decision to publish or in the preparation of the manuscript. The authors do not have any conflicts of interest to report.

## Notes

### Competing Interest Statement

The authors have declared no competing interest.

